# Evaluation of a deuterated triarylmethyl spin probe for in vivo R2^*^-based EPR oximetric imaging with enhanced dynamic range

**DOI:** 10.1101/2023.05.18.541366

**Authors:** Shun Kishimoto, Nallathamby Devasahayam, Gadisetti VR Chandramouli, Ramachandran Murugesan, Yasunori Otowa, Kota Yamashita, Kazutoshi Yamamoto, Jeffrey R Brender, Murali C Krishna

## Abstract

**Purpose:** In this study, we compared two triarylmethyl (TAM) spin probes, Ox071 and Ox063 for their efficacy in measuring tissue oxygen concentration by R2*-based oximetry in hypoxic and normoxic conditions.

**Methods:** The R2* dependence with spin probe and oxygen was calibrated using standard phantom solutions at 1, 2, 5, 10 mM spin probe and 0, 2, 5, 10, 21% oxygen concentrations. For a hypoxic model, in vivo imaging of a MIA PaCa-2 tumor implanted in the hind leg of a mouse was performed on successive days by using either Ox071 or Ox063. For normoxic model, renal imaging of healthy athymic mice was performed under similar conditions. The 3D spin density images acquired under three different gradients and reconstructed by single point imaging modality were used for computing the R2 * values.

**Results:** The signal intensities of Ox071 were about three times greater in the phantom solutions than Ox063 in the entire pO_2_ range investigated. Although histograms computed from the pO_2_ images of the tumor were skewed towards low pO_2_ levels for both spin probes due to R2* signal loss, more frequency counts in the normoxic region at pO_2_ > 32 mmHg could be detected with Ox071. In the normoxic model in kidney, the high pO_2_ cortex and the low pO_2_ medulla regions were well delineated. The histograms of high-resolution kidney oximetry images using Ox071 were nearly symmetrical and frequency counts were seen up to 55 mmHg which were missed in Ox063 imaging.

**Conclusion:** This study illustrates Ox071 as a better oximetric probe than Ox063 in terms of sensitivity and the pO_2_ dynamic range.

## 1. INTRODUCTION

Hypoxia, a critical factor in the progression and metastasis of many cancers, is strongly correlated with a poor prognosis. The outcome of therapeutical interventions, both of radiotherapy and chemotherapy, also depends upon the pO_2_ status of the tumor. These factors have led to the development of various techniques for determining the oxygenation status of tumors.^1-3^ Electron paramagnetic resonance (EPR)-based techniques offer high sensitivity for pO_2_ measurement of hypoxic tumors, because of the inverse relationship between the EPR signal intensity and the oxygen concentration. With the advantage of versatile and well-studied nitroxyl spin probes as well relatively simple instrumentation, the continuous wave (CW) EPR techniques have been on the forefront in the development and application of EPR oximetry.^4-6^ However, the advantages of time domain (TD) EPR, especially its temporal resolution for dynamic measurements, has been the driving force for addressing the challenges in the development of pulsed EPR instrumentation for in vivo studies.^7^ The development of single narrow EPR line spin probes, triarylmethyl (TAM) radicals, has enhanced the potential sensitivity and temporal resolution of pulsed EPR techniques for oxygen measurement.^8^ This class of spin probes has good chemical stability and long relaxation times. The EPR signals of TAM radicals are not generally broadened by binding with proteins and other biologic molecules, making them ideal sensors for TD EPR oximetry. Spin exchange between the TAM spin probe and the oxygen molecules decreases the TAM phase memory time, which could be readily measured by time domain EPR techniques. In addition, there is practically no self-broadening of the resonance in the concentration range of the spin probe used for imaging. The line broadening is nearly entirely dependent on the oxygen concentration, enhancing the selectivity of EPR oximetry.

TD EPR imaging using the TAM radicals offers fast acquisition of pO_2_ maps in vivo.^8-9^ The challenge of oximetry probe design is not only sensitivity and resolution but also the dynamic range of measurement. The median pO_2_ varies as high as 32 mmHg in tumors such as rectal carcinoma or to as low as 2 mmHg in some prostate and pancreatic cancers.^10^ Further, oxygen gradients develop in tumors as they grow. These hypoxic gradients play an important role in tumor migration.^11^ Hence accurately measuring pO_2_ in a large dynamic range of dissolved oxygen concentration with good linearity is of importance in cancer research. Such measurements will also mitigate the difficulty in the assessment of oxygenation of tumors prior to therapy as well as quantification of transient hypoxia. Organs such as kidney also have a range of pO_2_ distribution. The pO_2_ in renal cortical tissue is in the range of 20–60 mmHg. But the pO_2_ values of outer and inner medulla are in the range of 15–30 mmHg and < 15 mmHg, respectively.^12, 13^ BOLD-MRI has been used to detect R2 value changes in the cortex and medulla.^14^ In the renal cortex where pO_2_ is higher, most of the hemoglobin is in oxygenated state. The changes in pO_2_ in higher range (renal cortex) may not cause a large amount of the oxyhemoglobin dissociation, limiting the sensitivity of BOLD-MRI to detect changes in high pO_2_ in region.^13^ An oximetric probe with improved sensitivity and dynamic range for pO_2_ measurement will widen the applications of EPR oximetry.

Using the TAM spin probe Ox063, we have developed and assessed single point imaging (SPI) modality for pO_2_ measurement in vivo.^15-18^ Ever since the early report of TAM radicals, there have been continuous attempts to improve this family of EPR probes by structural modification as well as isotope enrichment. Ox071 is designed to make the resonance line narrower than that of Ox063, by replacing protons with deuterium nuclei as indicated in Figure1.^19, 20^ In a preceding study, we have shown that Ox071 spin probes can be used for pO_2_ estimation by measuring either R1 or R2*. Although R1-based pO_2_ estimation provided relatively better resolution of pO_2_ in standard solutions equilibrated with 0, 2, 5% oxygen, it required longer scan time which resulted in higher energy absorption.^21^ R2*-based oximetry can reduce the spectral as well as image acquisition time. On the other hand, pO_2_ images from R2*-based oximetry are prone to be noisy, especially at high delay times (t_p_ values) succeeding the radio frequency pulse. Our earlier observation that the R2* values of Ox071 were 3 times smaller than that of Ox063, in a wide range of oxygen concentration was suggestive to investigate the potential of Ox071 as a TD EPR oximetry probe. Here we present first 3D *in vivo* SPI oximetry using Ox071 spin probe. We compare the spin probes, Ox063 and Ox071 for their efficacy as oximetric probes for imaging hypoxic and normoxic regions by using a murine tumor model and healthy kidney in vivo.

**Figure 1.**
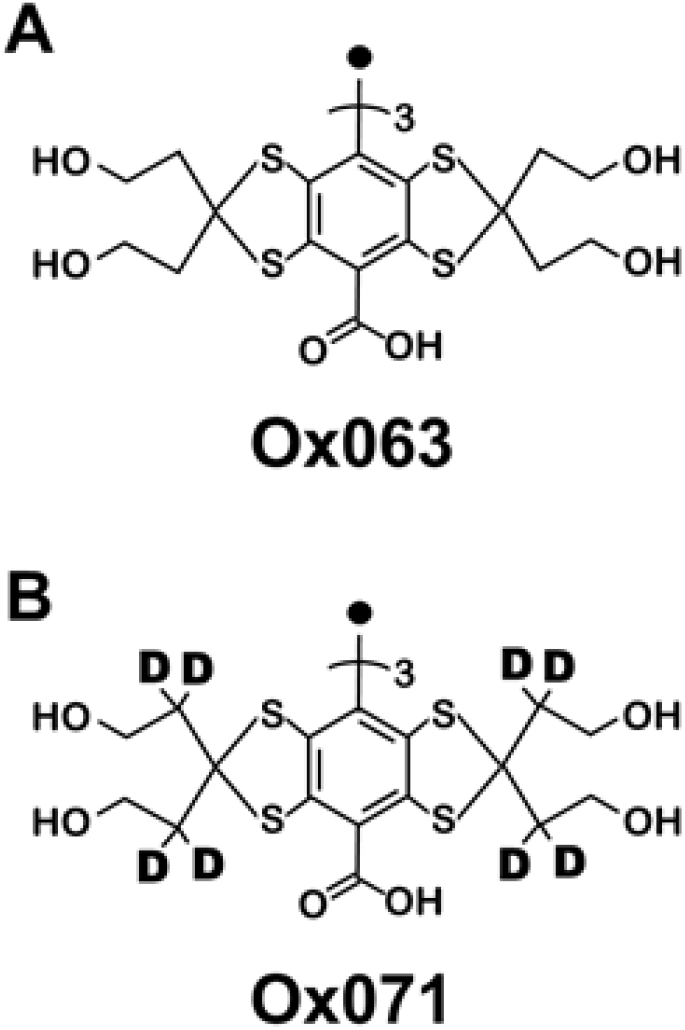
Schematic diagrams of the structures of Ox063 (A) and Ox071 (B). Ox071 is a deuterated version of Ox063 at methylene groups adjacent to thioketals.

## 2. METHODS

### 2.1. Animals

Female athymic NCr-nu/nu mice, supplied by the Frederick Cancer Research Center’s Animal Production unit (Frederick, MD), were housed in a climate-controlled and circadian rhythm-adjusted room and allowed food and water *ad libitum*. MIA PaCa-2 cells were obtained from ATCC (Manassas, VA 20110, USA) and the MIA PaCa-2 tumors of about 10 mm (diameter) were grown in the hind legs of 3 months old mice by injecting 3 × 10^6^ cells. Experiments were conducted in compliance with the Guide for the Care and Use of Laboratory Animal Resources (National Research Council, 1996) and were approved by the National Cancer Institute’s Animal Care and Use Committee.

### 2.2 Spectral data acquisition

A home-built time-domain EPR imaging system operating at 300 MHz was used for spectral and image data acquisition (9). The pO_2_ calibration experiments were performed using 2 mM Ox071 aqueous solutions filled in glass tubes and equilibrated at 5 different oxygen concentrations (0%, 2%, 5%, 10%, and 21%). Oxygen levels were achieved by bubbling appropriate O_2_ and N_2_ gas mixtures or argon gas through the sample for about 45 min. The spin probe concentration calibration experiments were done using 1, 2, 5, and 10 mM Ox071 solutions filled in separate glass tubes and equilibrated at 0% oxygen. Time-domain EPR signals were recorded by π/2-delay pulse sequence at a repetition time (TR) of 8 or 40 μs at ambient temperature. Spectral data acquisition parameters were: dwell time = 5 ns, number of averages = 40,000, number of phase cycles = 4, Number of samples = 640 and 2000 for TR = 8 and 40 μs, respectively. The R_2_* values were estimated by fitting the exponential decay of FID signal. The standard errors of R_2_* were estimated using 5 repeated FID signals.

### 2.3 Animal imaging

Mice, housed in a custom-designed holder, and maintained at a breathing rate of 60-80 per min and core body temperature of 37±1°C by a warm air blower, were anesthetized using 2-2.5% isoflurane in medical air. The rectal temperature was monitored by a nonmagnetic probe (FISO technologies, Quebec, Canada). A 30-G needle attached to polyethylene tubing was cannulated into the tail vein. In tumor imaging study, the hind leg of MIA PaCa-2 tumor bearing mouse was positioned in a 19-mm diameter parallel -coil resonator. For renal imaging of healthy animal, the mouse body was positioned in a 25-mm diameter parallel-coil resonator. A bolus (1.125 μM/g body weight) of 75 mM spin probe (Ox071/Ox063) solution was administered through tail vein cannulation.

### 2.4 Image acquisition and reconstruction

The anatomic map of mouse tumor was scanned using ICON 1T MRI scanner (Bruker Biospin) at matrix size = 128 × 128 × 14, field of view (FOV) = 28 mm and slice thickness = 2 mm. Images were acquired in the XZ-plane, which corresponds to the sagittal plane on the mouse leg. For comparison between Ox063 and Ox071, FT-EPRI scans were performed using the same mouse on successive days in tumor imaging study. For R_2*_ oximetry by SPI, the images were collected under three different maximum magnetic gradient values of 14, 11.4 and 9.6 mT/m. Mouse 3D imaging data were acquired at 19 × 19 × 19 Cartesian grid and the image reconstruction was done at 96 × 96 × 96 grid by zero-filling the k-space matrix. Data acquisition parameters were: dwell time = 5 ns, number of samples = 640, number of phase cycles = 4, number of averages = 4,000, TR = 8 μs and 3D image acquisition time = 12 min.

R_2_* maps were computed using 12 different t_p_ values from the multi-gradient data. The FOV and SNR are inversely proportional to the delay time t_p._ Hence the R_2_* map was computed by zooming all the images to equal FOV and intensity scaling to account for zooming. Image reconstruction was performed after correction for DC shifts and SNR of k-space by applying Tukey window (r = 0.7). Tissue pO_2_ images were computed from the R_2_* images using the calibration data. A high SNR intensity image was calculated by taking median of the 12 images and it was used to identify the background regions. The background regions of pO_2_ and S_0_ maps were masked for better visualization. The tumors were reconstructed at FOV ∼30 mm and co-registered with anatomy to locate the tumor regions. The kidney images were reconstructed at two different FOVs ∼30 and ∼24 mm to observe the shape and cross-section of kidneys in the intensity/spin density maps. These regions of interest (ROIs) were marked and masks were applied, based on ROIs to display pO_2_ values of kidney regions.

## 3. RESULTS

### 3.1 Estimation of calibration parameters

The principle of EPR oximetry relies on the Heisenberg exchange between an exogenous paramagnetic probe and the oxygen molecule which affects the relaxation of the spin probe. The relaxation rate R2* can be estimated from the free induction decay (FID) signal intensity S(t_p_) given by,

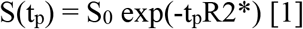

where S_0_ is the signal at t_p_ = 0. R2* and S_0_ were calculated from the slope and intercept of log(S(t_p_)) *vs*. t_p_ plot in the t_p_ range of 0.55 – 1.8 μs. For the TAM radical Ox063, the R2* showed a linear dependence with its concentration as well as with pO_2_ levels as reported earlier.^9^ For the spin probe Ox071, the concentration dependence of R2* was assessed by spectral measurements of 1, 2, 5, 10 mM of solutions equilibrated at 0% oxygen at a repetition rate of TR = 8 μs. The linear relationship (Figure. 2A) is given by,

**Figure 2.**
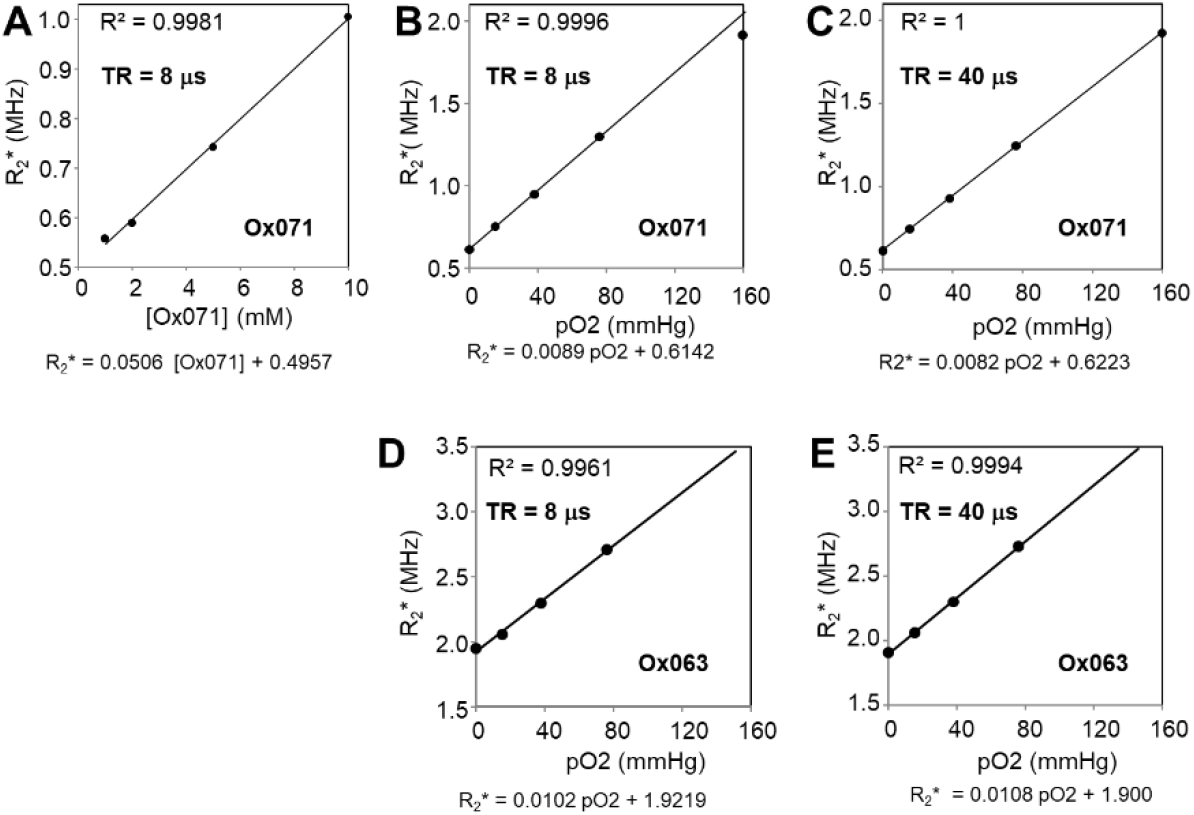
A – C. Linear relationships of Ox071. A. between R2* and Ox071 concentration for TR = 8 μs; B. between R2* and pO2 at TR = 8 (B), and between R2* and pO2 at TR =40 μs (C); The R2* is linearly proportional to pO2 in the range of 0 – 76 mmHg at TR = 8 μs which is routinely used for mouse experiments. R2* is linearly proportional to pO2 in the range of 0 – 159.6 mmHg at TR = 40 μs. D – E. Linear relationships of Ox063 between pO2 and R2* at TR = 8 (D) and TR = 40 (E).

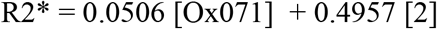

where R2* is in MHz and Ox071 concentration is in mM. The intercept in Eq. [2] corresponds to the natural linewidth of the Ox071. The repetition rate of the RF pulses, can be set at TR > 5T_1_ to allow the relaxation of excited spins to return back to ground state to give maximum FID intensity. However, for in vivo measurements, a short TR of 8 μs was selected due to scan time constraints originating from conditions such as the pharmacokinetics of the TAM radical or enhanced temporal resolution. At this TR, steady state free precession signals showed reasonably good SNR.

The relationship between R2* and pO_2_ was determined using 0, 2, 5, 10% and 21% oxygen-equilibrated Ox071 solutions (2 mM) at TR values of 4, 6, 8, 12, 15, 20, 28 and 40 μs. The R2* linearly increased with pO_2_ levels in the range of 0 – 10% for the entire range of TR studied. However, their slopes and intercepts were slightly different. The R2* vs. pO_2_ graphs at TR = 8 and 40 μs are shown in Figure 2B and Figure 2C, respectively. The relationships obtained for TR = 8 μs and TR = 40 μs are given in Eq. [3] and Eq. [4], respectively.

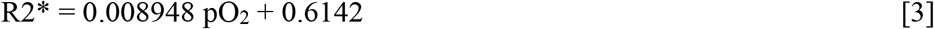

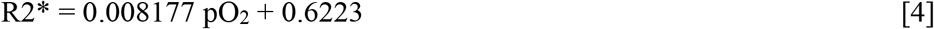

where R2* is in MHz and pO_2_ is mmHg. R2* vs. pO_2_ relationship for the spin probe, Ox063 evaluated at TR = 8 μs and TR = 40 μs (Figure 2D and 2E) is linear as given in Eq. [5] and Eq. [6], respectively.

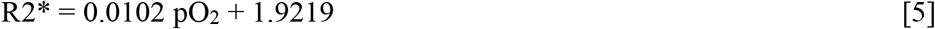

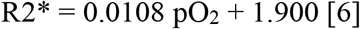

The intercepts in Eq. [4] and Eq. [6] indicate minimum R2* in the absence of oxygen for a 2 mM solution at TR = 40 μs. The minimum R2* of Ox063 (1.90 MHz) in eq. [6] is ∼3 times higher than that of Ox071 (0.62 MHz) in Eq. [4]. Thus as compared with Ox071, the FID of Ox063 decays faster and SNR decreases with time as shown in Figure. 3.

**Figure 3.**
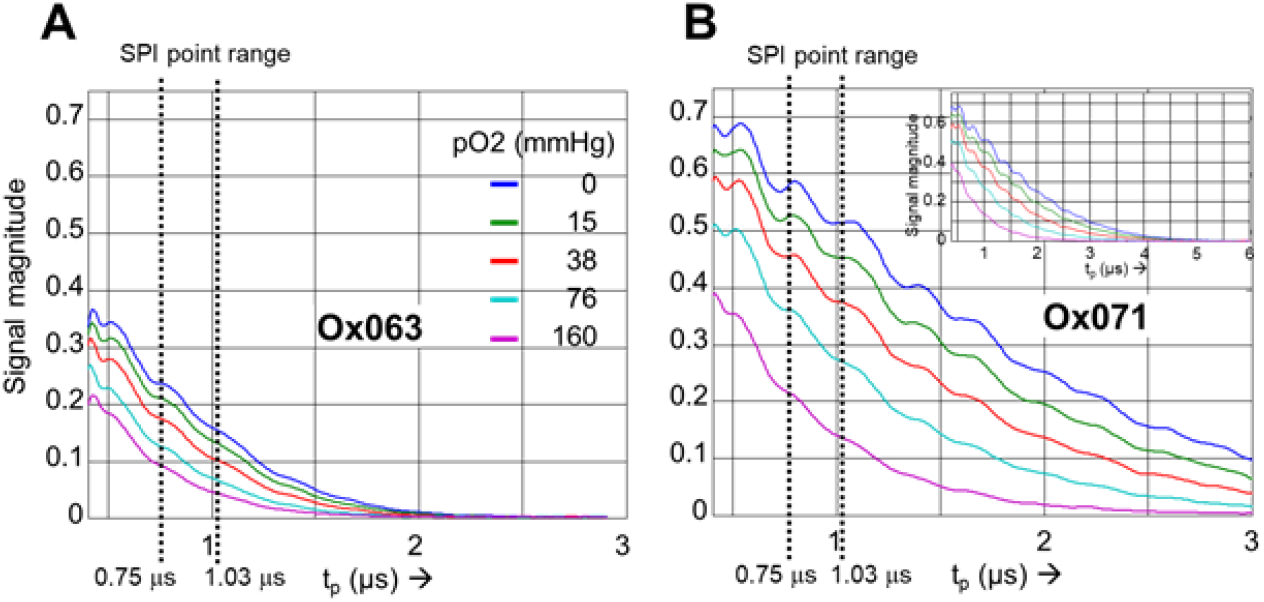
A. Signal decay of Ox063 at pO2 = 0, 7.6, 38, 76 and 159.6 mmHg in the tp range of 0.4 – 3.0 μs. B. Signal decay of Ox071 at the same pO2 values. Inset shows the decay in the range 0.4 – 6.0 μs. Signals were normalized to S0 = 1 by extrapolating the signal from the tp range of 0.5 – 1.5 μs to tp = 0. TR = 40 μs. Vertical dashed lines indicate tp range used for image reconstruction.

In SPI, the image is reconstructed at a given t_p_. The S_0_ and R2* maps are calculated from the intensities of a set of images obtained at several t_p_ values. The images at different t_p_ values are scaled to equal FOV, intensities are corrected to FOV scaling and S_0_ and R2* are calculated using the Eq. [1]. The fitting procedure leads to higher noise in both S_0_ and R2* maps at low intensity regions such as in the background. The t_p_ range used for routine mouse oximetry is shown by dashed vertical lines in Figure 3. A comparison of Ox063 and Ox071 profiles indicates that the signal magnitudes of Ox071 are nearly twice to that of Ox063 in the selected t_p_ range. The SNR and FOV decrease as the t_p_ is increased. For large t_p_, the resolution is higher, but the poor SNR can make the images noisy, compromising the accuracy of the pO2 maps. In this context, Ox071 is a better spin probe than Oxo63, and capable of providing high resolution images at better SNR.

### 3.2 Mouse tumor imaging

MIA PaCa-2 tumor, implanted in a mouse hind leg was imaged using both the spin probes, Ox071 and Ox063, on successive days. One day interval was set between Ox071 and Ox063 imaging for clearance of Ox071 from the tumor.^22^ The zero gradient signal intensities from these imaging experiments are compared in Figure 4B. The signal level of Ox071 is approximately twice stronger and lasts twice longer as compared to that of Ox063. The R2* map was calculated from 12 delay time points starting from 0.75 μs (marked by the vertical dashed line in Figure 4B) at 30 ns intervals. The intensities of the voxels were fitted to Eq. [1] to obtain spin density (S_0_) and pO2 maps. The cross sections of S_0_ and pO2 maps of the tumor at a selected slice (61^st^ of 96) are shown in Figure 4C (the tumor is indicated by a dashed gray line). The spin density maps of Ox071 and Ox063 show about 6 high intensity regions distributed across the tumor, but these were more diffuse in Ox063 than in Ox071. The uneven distribution of the probe is probably due to the presence of poorly developed vasculature within the tumor region where the perfusion of probe is relatively less. The pO2 maps assessed by Ox063 and Ox071 also appear to be similar to each other by displaying five high pO2 regions distributed circularly within the tumor. A difference between Ox063 and Ox071 maps may be seen at top right. It is where the bladder is located, and the difference is likely due to the differences in accumulation of the spin probe in bladder during the experiment. Such probe accumulation in the bladder is generally observed during EPR imaging.

**Figure 4.**
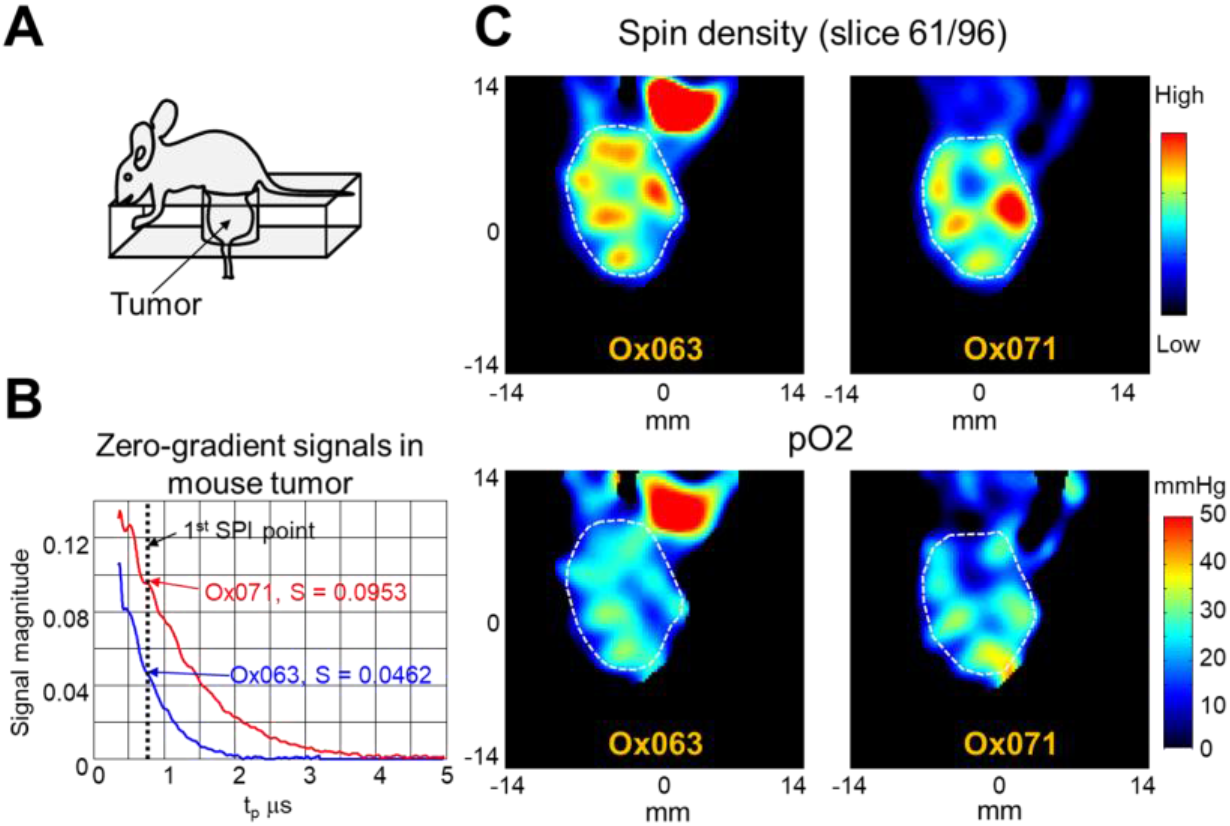
Tumor imaging by the spin probes Ox063 and Ox071. A. Mouse tumor position in resonator. B. Zero-gradient signal intensities of Ox063 (blue) and Ox071 (red) observed in mouse tumor. Vertical dashed line indicates a delay time t_p_ used for image reconstruction. Note the signal level of Ox071 is about double to Ox063 at this point. C. Spin density (top) and pO_2_ (bottom) maps of a slice (61/96) through the tumor bulk determined by the probes Ox063 (left) and Ox071 (right). Tumor region is enclosed by dashed gray lines.

EPR imaging is a functional imaging modality and it provides the visualization of the spin probe distribution. To relate to the tumor anatomy, MR imaging was performed with a matrix size of 128 ×128 × 14. A slice covering the tumor region (slice 9/14) is shown in Figure. 5A. The EPR 3D spin density and pO2 maps were co-registered with anatomic maps from MRI. The MRI was used as base map and 3D EPR data was transformed by translation, rotation and scaling operations and finally the transformed data was used to calculate EPR data slices that are geometrically equivalent to MRI slices. The tumor regions were marked based on the MR images. The overlay of pO2 on an anatomic slice is shown in Figure 5B and 5C. The pO2 values of tumor region in all slices were determined, and pO2 histograms of tumor region were calculated for Ox063 (Figure 5D) and for Ox071 (Figure 5E). The pO2 histogram of Ox071 is more symmetric than that of Ox063. Ox063 histogram is left-skewed and there is almost nil frequency distribution above pO2 > 30 mmHg. A closer look at the histograms revealed that frequencies for pO2 > 30 mmHg are well seen in the case of Oxo71 (Figure 5G) as compared to Ox063 (Figure 5F).

**Figure 5.**
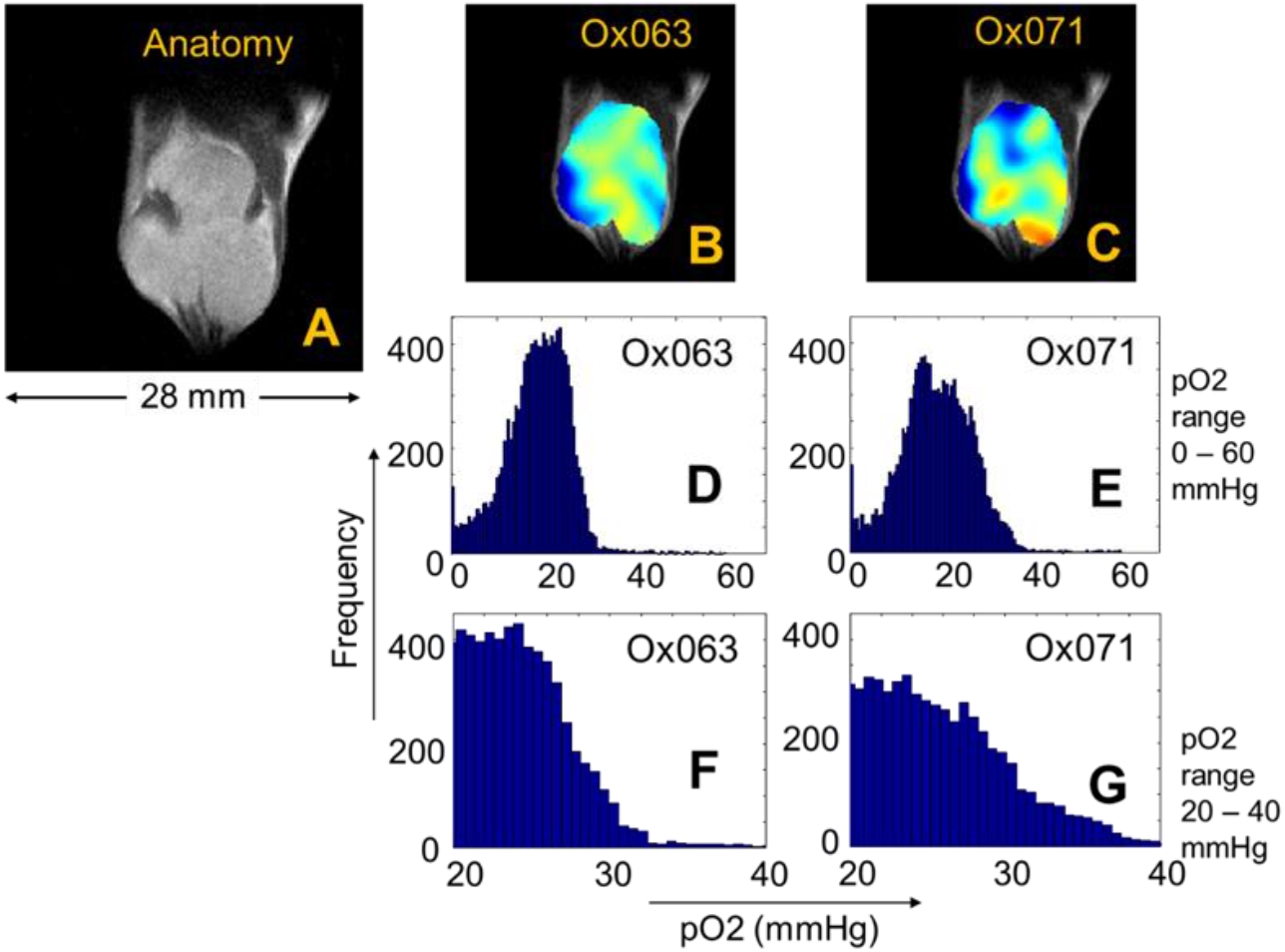
Histograms of voxel pO2 values in tumor. A. MRI. Anatomic map of a slice (9 of 14) through tumor bulk. B and C. Co registration of MRI (A) and pO2 map obtained using Ox063 (B) and Ox071 (C). D and E. Histograms of pO2 values of tumor voxels determined by Ox063 (D) and Ox071(E) respectively. F and G. Distribution of pO2 values obtained by Ox063 and Ox071 in the range of 20 – 40 mmHg. Note higher frequency counts of Ox071 in G at pO2 >30.

### 3.3 Renal imaging

To further assess the large dynamic range (especially higher pO2 region) capability of Ox071 in TD EPR oximetry, we performed renal imaging of healthy mice using both Ox071 and Ox063 probes. The zero gradient signal intensities from both imaging data were compared in Figure 6A which showed definite advantage of Ox071 above t_p_ > 0.7 μs. Both Ox063 and Ox071 probes remained in kidneys for longer period of time compared to any other solid organs. Spin density images of one of the slices of the 3D images of the kidney using Ox063 and Ox071 were shown in Figure 6B. The SNR of the spin density image of Ox071 imaging was higher than that of Ox063 imaging. The corresponding pO2 images derived from R2 * values were also presented in Figure 6B. The pO2 level was clearly higher in cortex region than medulla/renal pelvis region in both images. However, spatial resolution of 1.6 mm was not high enough for clear distinction.^23, 24^ To evaluate the advantage of the narrow resonance of Ox071, the pO2 image was constructed using higher t_p_ value to enhance the spatial resolution to 1.2 mm. Figure 6D shows the comparison between Ox071 and Ox063 kidney-oximetry at higher resolution. Interestingly, both probes distinguished cortex from medulla/renal pelvis. However, in the case of Ox071, the pO2 map indicates the cortex region is well oxygenated than that of Ox063 image. The histogram of the high resolution pO2 images, presented in Figure 7A, showed that the Ox071 was capable of imaging oxygen concentration even a little above 50 mmHg. On the other hand, Ox063 showed relatively lower pO2 range. The median (± SD) pO2 values estimated by Ox071 and Ox063, respectively are 40.5 (±4.7) and 35.6 (±3.9) and the difference was found to be statistically significant (p = 0.04).

**Figure 6.**
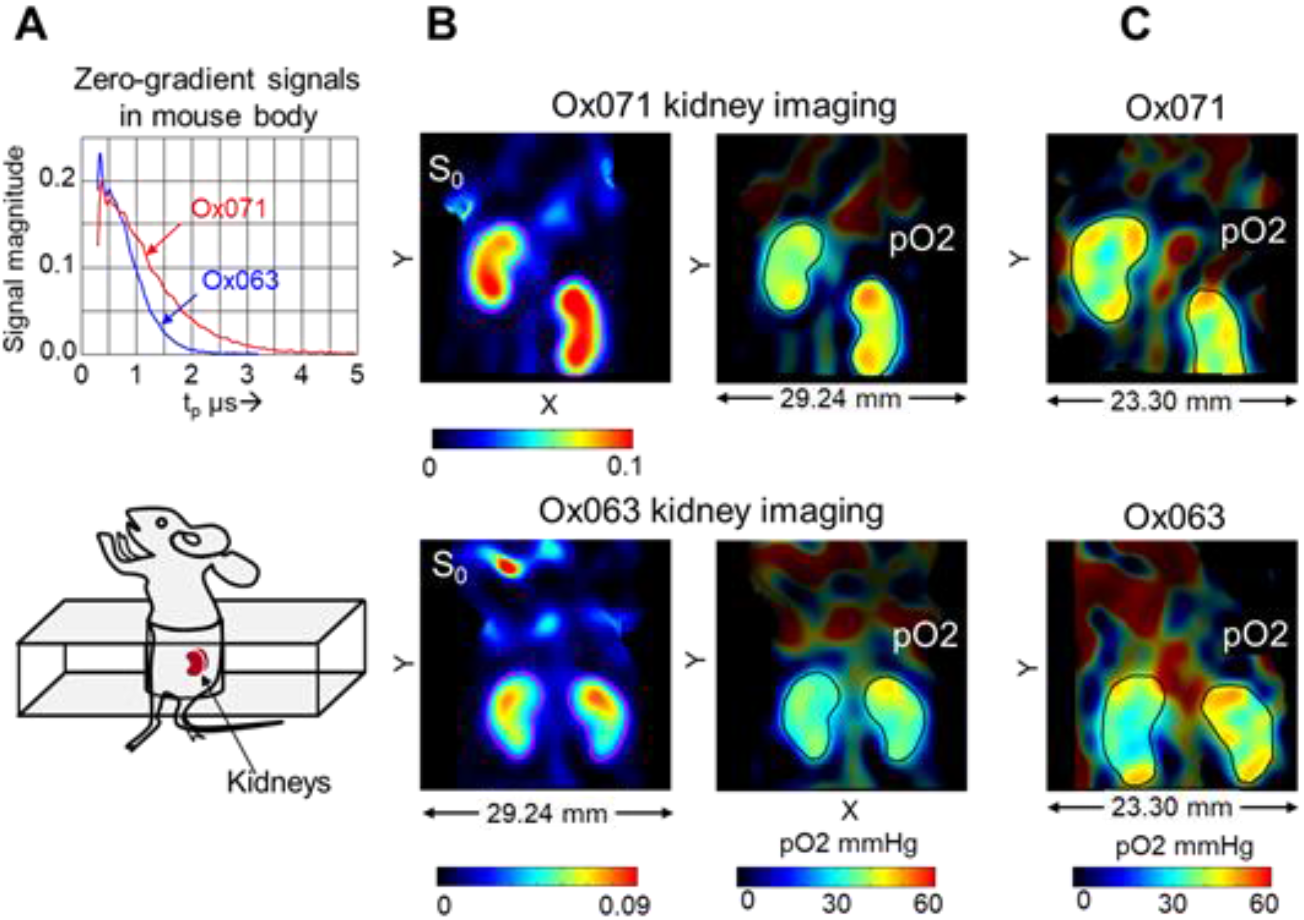
A. Zero-gradient signal intensities of Ox063 (blue) and Ox071 (red) observed in whole body imaging of two different mice. B. The spin density (S_0_) and pO_2_ maps of a slice in kidney region at 1.6 mm spatial resolution. Top row: Imaging using Ox071 showing a slice (75/96) across kidneys. Bottom row: Imaging using Ox063 in a different mouse (slice 74/96). The kidney regions are enclosed by ROI lines both in S_0_ and pO_2_ maps. The pO_2_ values outside ROIs are shown at brightness relative to the probe abundance. C. The kidney pO_2_ maps calculated at higher spatial resolution (1.2 mm) using Ox071 (slice 82/96, top) and Ox063 (slice 82/96, bottom).

**Figure 7.**
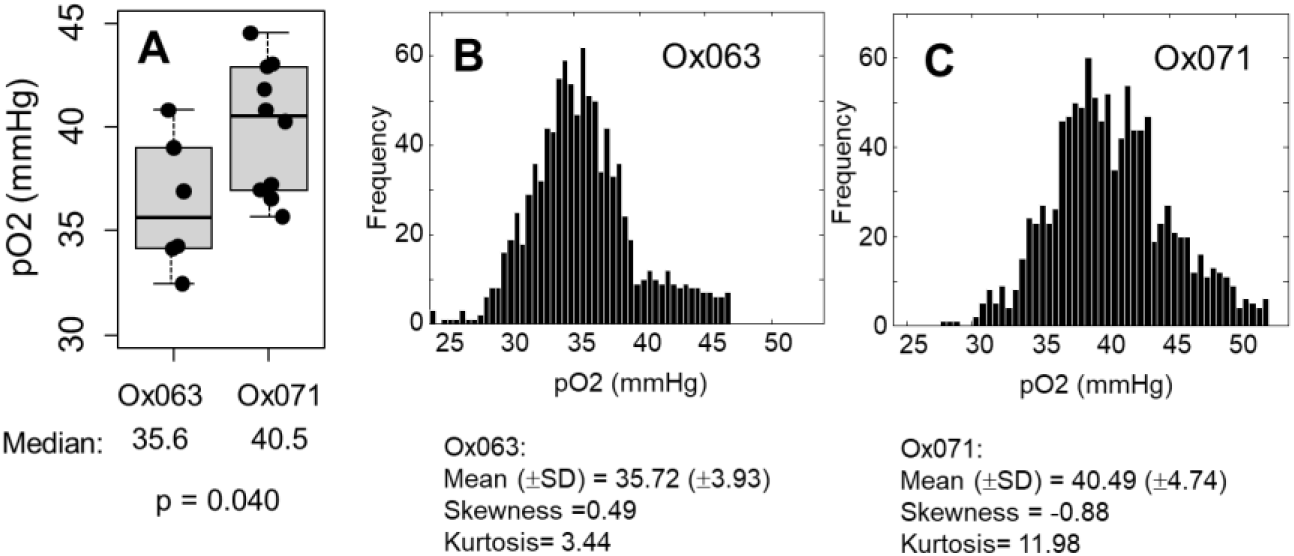
A. Box plot of median pO2 values in kidney regions determined by Ox063 (n = 6, 3 mice) and Ox071 (n = 10, 5 mice). The thick line inside the box is median, hinges are 1st and 3rd quartiles and notches extend till 1.5 IQR. B & C. Histogram of voxel pO2 values in kidney determined by Ox063 (B) and Ox071 (C).

## 4. DISCUSSION

Repeated quantitative measurement of tumor hypoxia may be useful to evaluate tumor treatment and prognosis. EPR oximetry has evolved as a viable tool for in vivo pO2 measurement. To enhance its potential, new technologies and methodologies are under development in many laboratories. The significant advancement made in time domain EPR instrumentation has enabled to measure tissue oxygen status in vivo in tumor models with good temporal and spatial resolution. However, translation of this capability for useful preclinical applications is faced with some constraints and challenges in terms of suitable time domain imaging methodologies and spin probes with good sensitivity and linearity for pO2 measurement in a wide range of pO2 values. Triarylmethyl radicals (TAMs) have been the ideal choice for TD EPR oximetry because of their extraordinary in vivo stability, very narrow single resonance and enhanced sensitivity to O_2_. Using the TAM radical Ox071 as a spin probe we have evaluated R1 and R2*-based pO2 images from TD EPR imaging. SPI reconstruction resulted in noisy images at higher pO2 regions because of the inverse dependence of the phase memory time with oxygen concentration.

The present study aimed to evaluate Ox071, a spin probe with narrower linewidth than that of Ox063, for its potential in in vivo TD EPR oximetry. Compared to Ox063, Ox071 showed high intense FIDs with long decay time (Figure 3). For example, the signal intensity of 2% oxygen saturated 2 mM Ox071 solution which corresponds to hypoxic tissue pO2 levels showed 2.3 times stronger signal than that of Ox063 at the delay time t_p_ = 0.75 μs. The difference increased to a factor of 2.6 at 5% oxygen which corresponds to normal tissue oxygen concentration. At a delay time of 1.03 μs the difference increased by a factor of ∼3.3 and ∼3.7, respectively for 2% and 5% oxygen concentrations. The data suggest that Ox071 will provide higher signal intensity in most of the organs even at high oxygen levels, resulting in superior estimates of R2*.

TAM-based spin probes found application in tumor oximetry, because their clearance in tumors is slow due to the enhanced permeability and retention effect, arising from the tumor vasculature. Mapping pO2 distribution in tumors can identify hypoxic fractions that are known to be less sensitive to therapeutic interventions. The results of MIA PaCa-2 tumor imaging given in Figure 4 present a comparative evaluation of Ox063 and Ox071 for tumor imaging. The zero-gradient FID profiles of Ox063 and Ox071 in the tumor in vivo were similar to the observations made in phantoms where Ox071 showed stronger and longer FIDs. The distribution of both probes in the tumor looked nearly the same as seen in the spin density images (Figure 4C top row). Histogram representations of the tumor pO2 data, computed from R2* images are given in Figure 5 D-G. The form of the histogram can provide information about the heterogeneity of tumor oxygen status. The pO2-histograms are expected to be bell-shaped, following a Gaussian distribution in a homogeneous phase. However, most tumors are reported to have a skewed distribution because of their hypoxic nature. The skewness depends on the degree of hypoxia. The first look at the histograms of Ox063 and Ox071 (Figure 5D and 5E) may indicate that both of them are similar. However, the expanded versions given in Figures 5F and 5G reveal that Ox063 has missed frequency counts at pO2 > 32 mmHg. The R2* values of Ox063 was nearly three times higher than that of Ox071 at this pO2 (Figure 2). Consequently, the loss of signal intensity is higher for Ox063 which may have resulted in missing the information at higher pO2 region. The tumor voxel pO2 mean (±SD), skewness and kurtosis estimated by Ox071 are: 19.6 (±8.5) mmHg, 0.27 and 4.1 and estimated by Ox063 are: 19.0 (±7.2) mmHg, -0.12 and 4.3 respectively.

The superior performance of Ox071 in terms of enhanced dynamic range was more evident in kidney oximetry. Kidney pO2 images, obtained using both the spin probes showed higher oxygen concentration at the cortex region. Healthy kidneys are known to have pO2 distribution ranging from 20 mmHg at medulla region to 60 mmHg at cortex region. The high-resolution kidney oximetry using Ox071 showed pO2 gradient between high pO2 cortex region and lower pO2 medulla/renal pelvis region. The histograms of high resolution pO2 images (Figure 7B) brought out the difference in the dynamic range capability of the probes. Ox071 histogram is nearly symmetrical reflecting homogeneous nature of spin probe distribution in the highly vascular kidneys. On the other hand, Ox063 histogram is right skewed. While frequency counts are seen up to 55 mmHg in Ox071 imaging, the probe Ox063 misses the high pO_2_ region. In the previous studies of pO_2_ measurement using electrode, it is known that the pO_2_ values in cortex region are between 40-50 mmHg and in medulla/renal pelvis region are around 25 mmHg.^25-27^ The kidney slice voxel pO_2_ mean (±SD), skewness and kurtosis estimated by Ox071 are: 40.5 (±4.7) mmHg, -0.88 and 12.0 and estimated by Ox063 are: 35.70 (±3.9) mmHg, 0.49 and 3.4 respectively. The cortex and medulla regions in the pO_2_ images can be further resolved by increasing the image matrix dimension size from 19 to higher values and also using high t_p_ values in SPI reconstruction, because the steady-state spin probe concentration is quite amenable for the image collection time. The normoxic imaging experiment further validates the enhanced–dynamic range capability of the spin probe Ox071 in TD EPR oximetry.

The oxygenation status of kidney also provides important information for patients with chronic kidney disease (CKD). Renal hypoxia, particularly in the tubulointerstitium, is a candidate prognostic marker for CKD progression.^28^ Future high-resolution kidney oximetry using Ox071 may be useful to evaluate the oxygenation changes in medulla and cortex regions which may find application in studying kidney diseases such as renal artery stenosis. These studies as well as studies such as tumor oxygenation may benefit from the temporal resolution enhancement strategies available in TD EPR. In the present study data acquisition for a 3D image of 19x19x19 size requires 12 minutes. The temporal resolution can be enhanced by reducing the pulse repetition rate, TR. Using short TR values for spin probes with long relaxation time can result in interfering echoes. Thus short TR rates for Ox071 imaging may result formation shadows of low intensity images due to echo signals (TR).^21^ However, the shadows formed by echoes may be disregarded since the phase evolution of decaying signal is reverse to the echo signal. The delay times, t_p_ for which the echo intensities are appreciable vary with TR. The fitting procedure adopted using 12 delay points to calculate S_0_, and R2* treats them as deviations from fit line. Further, the delay points having no echo contributions can be identified *apriori* by measuring a phantom. A phantom containing 3 tubes filled with Ox071 solution was imaged to examine echo contributions. No echo interference was found in the t_p_ range used for mouse image formation.

## 5. CONCLUSION

This study has made a comparative evaluation of two TAM spin probes, Ox063 and Ox071 for their potential as oximetric probes in time domain EPR, by using pancreatic ductal adenocarcinoma xenograft MIA PaCa-2 tumor inoculated in the hind leg as a hypoxic organ, and healthy kidney as a normoxic organ in a mouse model. The pO_2_ images were computed using R2* values derived from three gradient SPI modality. The relatively narrower linewidth of Ox071 provided images with good signal intensity. The tumor pO_2_ histograms obtained using Ox063 were left skewed and had less frequency count in the high pO_2_ region. The renal imaging of healthy mice showed that the cortex regions can be resolved from the medulla in the pO_2_ maps. The high resolution kidney images from Ox071 probe were less noisy and the histograms showed that the high range pO_2_ values can be better captured by the Ox071, validating that oximetry using Ox071 has a wider dynamic range, compared to Ox063. The results point out to the importance of the knowledge of the spin probe characteristics before interpreting in vivo pO_2_ data. Tuning the sensitivity and dynamic range of TD EPR oximetry can widen its potential to study various pathophysiological conditions where oxygen concentration plays vital role.

## ACKNOWLEDGEMENTS

No competing financial interests exist. This research was supported by intramural funds from the Center for Cancer Research of the National Institutes of Health

## Notes

### Competing Interest Statement

The authors have declared no competing interest.

